# Comparative Performance of Popular Methods for Hybrid Detection using Genomic Data

**DOI:** 10.1101/2020.07.27.224022

**Authors:** Sungsik Kong, Laura S. Kubatko

## Abstract

Interspecific hybridization is an important evolutionary phenomenon that generates genetic variability in a population and fosters species diversity in nature. The availability of large genome scale datasets has revolutionized hybridization studies to shift from the examination of the presence or absence of hybrids in nature to the investigation of the genomic constitution of hybrids and their genome-specific evolutionary dynamics. Although a handful of methods have been proposed in an attempt to identify hybrids, accurate detection of hybridization from genomic data remains a challenging task. The available methods can be classified broadly as site pattern frequency based and population genetic clustering approaches, though the performance of the two classes of methods under different hybridization scenarios has not been extensively examined. Here, we use simulated data to comparatively evaluate the performance of four tools that are commonly used to infer hybridization events: the site pattern frequency based methods *HyDe* and the *D*-statistic (i.e., the ABBA-BABA test), and the population clustering approaches *structure* and ADMIXTURE. We consider single hybridization scenarios that vary in the time of hybridization and the amount of incomplete lineage sorting (ILS) for different proportions of parental contributions (*γ*); introgressive hybridization; multiple hybridization scenarios; and a mixture of ancestral and recent hybridization scenarios. We focus on the statistical power to detect hybridization, the false discovery rate (FDR) for the *D*-statistic and *HyDe*, and the accuracy of the estimates of *γ* as measured by the mean squared error for *HyDe, structure*, and ADMIXTURE. Both *HyDe* and the *D*-statistic demonstrate a high level of detection power in all scenarios except those with high ILS, although the *D*-statistic often has an unacceptably high FDR. The estimates of *γ* in *HyDe* are impressively robust and accurate whereas *structure* and ADMIXTURE sometimes fail to identify hybrids, particularly when the proportional parental contributions are asymmetric (i.e., when *γ* is close to 0). Moreover, the posterior distribution estimated using *structure* exhibits multimodality in many scenarios, making interpretation difficult. Our results provide guidance in selecting appropriate methods for identifying hybrid populations from genomic data.

---

While phenotypes of interspecific hybrids that are intermediate to two parental populations have historically been used as evidence of hybridization (e.g., Anderson, 1953; Hermansen et al., 2011; Isomura et al., 2013; Lehtinen et al., 2016; Pauers et al., 2018), the use of phenotypic data for identification of hybrids is often impractical because the range of such intermediacy can be overwhelmingly large (Anderson, 1953; Mallet, 2005). In addition, phenotypes can sometimes appear to be counterintuitive. For example, heterosis or transgressive segregation, where the phenotype of the hybrid falls outside of the range of parental variation, and adaptive introgression, where the hybrid closely resembles one of the parents via backcrossing (see review by Goulet et al., 2017), are cases in which hybrid detection would be challenging using phenotypic data alone.

The availability of ever-increasing amounts of genomic data coupled with advances in statistical and computational methods for handling large datasets has enabled researchers to use genomic data for detecting signals of hybrid ancestry in the past history of a species (Twyford and Ennos, 2012). While heterogeneity in the gene tree distribution across the genome of a hybrid achieved by genomic mosaicism (Folk et al., 2018) is very useful for this task, other evolutionary processes may result in similar genomic footprints, including incomplete lineage sorting (ILS), horizontal gene transfer, recombination, gene duplication and loss, and historical nonrandom mating, among others (Maddison, 1997). Much methodological work has been devoted to detecting hybridization in the presence of ILS as both processes are likely to occur simultaneously (Holder et al., 2001; Joly et al., 2009; Kubatko, 2009; Meng and Kubatko, 2009; Solís-Lemus and Ané, 2016). We briefly introduce several model-based methods that are popularly employed to detect hybridization.

Patterson’s *D*-statistic (Green et al., 2010; Durand et al., 2011; Patterson et al., 2012), popularly referred to as the ABBA-BABA test, is widely used to conduct a hypothesis test to infer introgression in spite of ILS. It is based on the magnitude of the observed difference in two discordant bi-allelic site pattern frequencies. The *D*-statistic is robust across a wide range of genetic distances although it may be sensitive to population size (Green et al., 2010; Jónsson et al., 2014; Martin et al., 2016; Kumar et al., 2017; Zheng and Janke, 2018). Refinements of the method continue to be explored in recent studies, including extensions that allow for application of the test to cases beyond the four-taxon phylogeny as well as those in which multiple individuals per population are sampled (Pease and Hahn, 2015; Elworth et al., 2018; Soraggi et al., 2018; Hibbins and Hahn, 2019; Malinsky, 2019).

Kubatko and Chifman (2019) proposed a method based on phylogenetic invariants for detecting both recent and ancestral hybridization events that considers both ILS and hybrid speciation within the same framework. Note that hybrid speciation differs from introgression as the taxon resulting from the process of hybrid speciation is maintained as a distinct taxonomic entity, whereas in the case of introgression, the hybrid entity merges with one of the parents via repeated backcrossing. The two processes will thus result in different topologies, as shown in Figure 1a (hybrid speciation) and 1b (introgression). Implemented in the computer program *HyDe* (Blischak et al., 2018), this method can examine genomic-scale data for a large number of taxa and detect the population(s) that may have arisen via hybrid speciation as well as their putative parental populations with statistical power that is similar to the *D*-statistic (Kubatko and Chifman, 2019). In addition to conducting a formal statistical test for hybridization, *HyDe* estimates the inheritance parameter, *γ*, that quantifies the genomic contributions of the parents to the hybrid and has been useful in detecting hybridization in frogs (Chan et al., 2019), Cichlid fish (Olave and Meyer, 2020), Persian Walnut (Zhang et al., 2019), and polyploid trees (Wagner et al., 2019). Because *HyDe* is a site pattern based method like the *D*-statistic, it is computationally efficient and not limited by the size of the dataset. However, genome-scale data may be necessary for optimal performance in complex hybridization scenarios (Blischak et al., 2018).

**Fig. 1.**
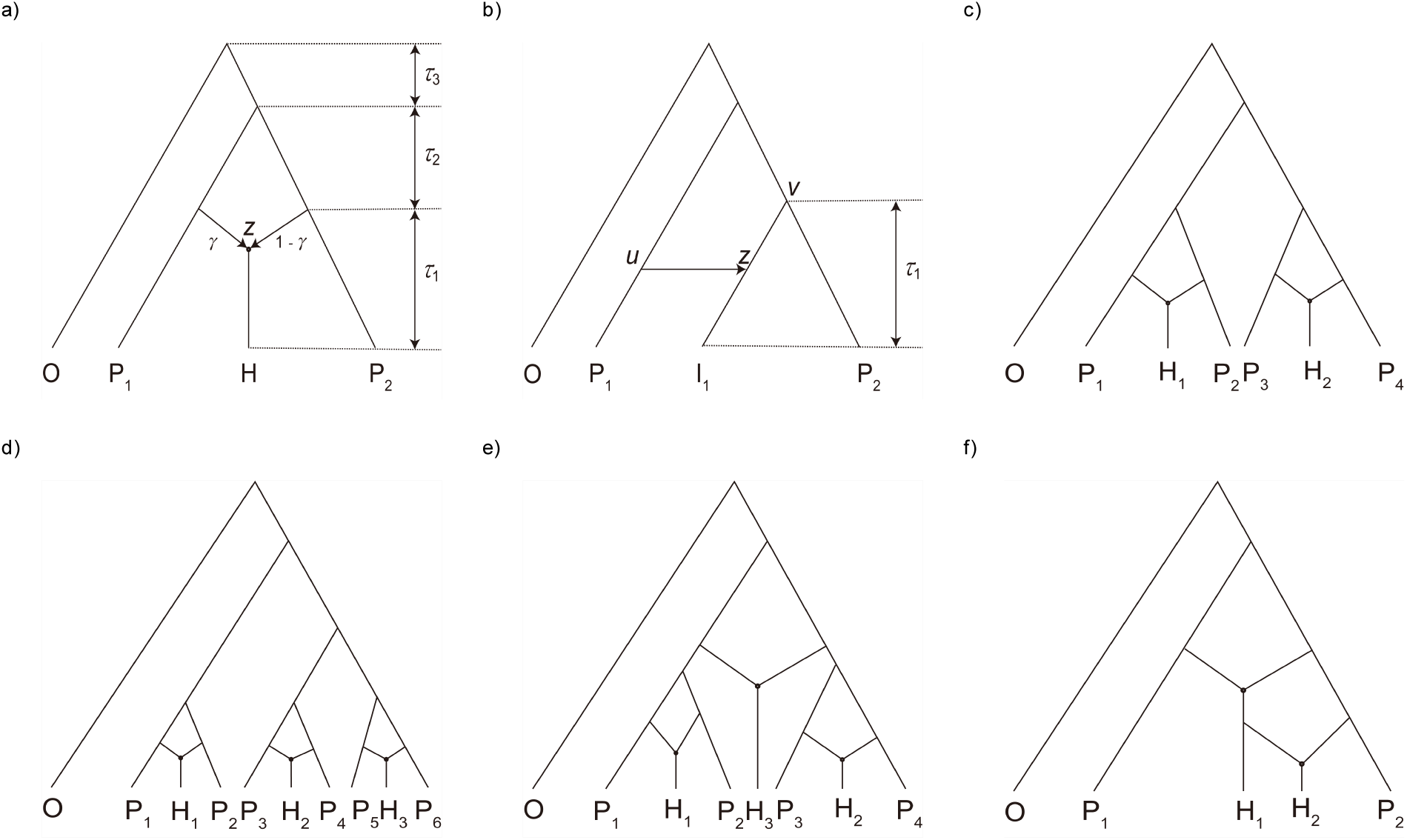
Six hybridization scenarios considered in this study: a) single hybridization event, b) introgression, c) two independent hybridization events, d) three independent hybridization events, e) three hybridization events that overlap (i.e., parental lineages are involved in more than one hybridization event), and f) mixture of ancestral and recent hybridization events. The outgroup population is labelled as O, parental populations are labelled as P_*i*_, and hybrids or introgressants are labelled as H_*i*_ or I_*i*_, respectively. Edge length, or the interval between speciation events, is denoted as *τ*_*i*_. Distinct populations and *τ* s are labelled with different integers in subscript *i*, and *τ*_1_ is more recent than *τ*_2_. In a), *γ* and 1 - *γ* below each hybrid edge denote the contribution of the corresponding parental population to the genome of the hybrid. The hybrid node is labelled as *z* in a) and b). In b), the two parental tree nodes that contribute to the hybrid node *z* are denoted by *u* and *v*.

Population genetic clustering approaches are popularly employed to identify hybrid individuals from genetic data and to hypothesize historical processes that shaped present population structure. Often implemented within a maximum likelihood (ML; e.g., ADMIXTURE; Alexander et al., 2009) or Bayesian-inference (BI; e.g., *structure*; Pritchard et al., 2000) framework, these methods provide a statistical basis to estimate the contribution of various genetic groups to an individual’s ancestry through estimation of probabilistic quantities called ancestry coefficients. These coefficients are interpreted as the proportion of genetic material inherited from ancestral gene pools to an individual in contemporary populations, but they are also often interpreted to be equivalent to the proportion of parental contributions in a hybrid population (Vähä and Primmer, 2005). There is no standardized and objective way to interpret these values, and most studies simply set an arbitrary probability threshold for use in classifying an individual as pure or admixed (Randi, 2008; Khosravi et al., 2013; Ito et al., 2015). Reliance on the ancestry coefficient can lead to mis- or overinterpretation of the processes in nature because different evolutionary scenarios can result in indistinguishable ancestry coefficients (Barilani et al., 2007; Anderson and Dunham, 2008; Lawson et al., 2018). Moreover, these methods are sensitive to the choice of markers, the level of genetic differentiation between populations, and the amount of data utilized (Vähä and Primmer, 2005; Latch et al., 2006; Kalinowski, 2011). Importantly, many of these methods were not originally designed to detect hybrid individuals, although they are popularly employed for this task.

The ability of the different population clustering methods to identify hybrids has previously been compared and evaluated. Two BI methods, *structure* and *NewHybrids* (Anderson and Thompson, 2002), were found to perform similarly well in detecting recent hybridization using simulated data with varying levels of genetic divergence and numbers of loci, although *NewHybrids* performed slightly better when the dataset contained a mix of F_1_ hybrids and introgressants (Vähä and Primmer, 2005; Oliveira et al., 2015). Comparisons of *structure, NewHybrids, BAPS* (Corander and Marttinen, 2006) and GeneClass (Piry et al., 2004) using simulated hybrids showed that the former three full BI methods outperformed the semi-Bayesian approach employed by GeneClass (Sanz et al., 2009). However, all methods showed reduced efficiency in their ability to detect hybrids as the rate of backcrossing increased. Bohling et al. (2013) and Neophytou (2014) compared the performance of *structure* and *BAPS* and found that *structure* was more likely to detect admixed populations and accurately estimate the ancestry coefficients, although misclassification occasionally occurred. Interestingly, both methods were outperformed by an ML-based assignment program (the Canid Assignment Test; Bohling et al., 2013).

In this study, we aim to comparatively evaluate the performance of four methods that are commonly used for hybrid detection using genomic data, *HyDe*, the *D*-statistic, *structure*, and ADMIXTURE, in a wide range of hybridization scenarios using simulated data. We consider six evolutionary scenarios: (1) a single hybridization event that varies in the time of hybridization and the amount of ILS for different values of *γ*; (2) introgression; (3) two and (4) three recent independent hybridization events; (5) three hybridization events that overlap (i.e., parental lineages are involved in more than one hybridization event); and (6) a mixture of ancestral and recent hybridization events. The evaluation is based on the power of the methods to detect hybridization and on the false discovery rates (FDR) for the *D*-statistic and *HyDe*, while the accuracy of estimated parental contributions is evaluated using mean squared error (MSE) for *HyDe, structure*, and ADMIXTURE.

## Materials and Methods

### Data Simulation

The six hybridization scenarios considered in this study are illustrated in Figure 1. For the single hybridization scenario (Fig. 1a), we used several different combinations of parameters: *τ*_1_ ∈ {0.05, 0.5, 1.5, 2.5, 5} with *τ*_2_ fixed at 1.5 (all branch lengths are measured in coalescent units, defined to be the number of 2*N* generations, where *N* is the effective population size) and *γ* ∈ {0, 0.1, 0.2, 0.3, 0.4, 0.5}, producing 30 combinations; and *τ*_1_ fixed at 1.5 with *τ*_2_ ∈ {0.05, 0.5, 2.5, 5} and *γ* ∈ {0, 0.1, 0.2, 0.3, 0.4, 0.5}, leading to another 24 combinations. Larger values of *τ*_1_ lead to increases in the amount of time that has elapsed since hybridization, which allows time for mutations to accumulate in the descendant lineages, while smaller values of *τ*_2_ lead to increases in the amount of ILS. We chose 1.5 for the ‘default’ edge length as it will result in a modest amount of ILS (Degnan and Rosenberg, 2006). The edge that links the root and the most recent common ancestor (MRCA) of the ingroup populations (*τ*_3_) was set to 5.0 to ensure sufficient genomic differentiation of the outgroup. For the scenarios in Figure 1b–1f, we assigned a single set of parameters: *γ* = 0.5 with all *τ* fixed to 1.5 except for the edge linked to the root and MRCA of the ingroup populations, which was set to 5.0.

For each condition, 10^3^ gene trees were simulated using *ms* (Hudson, 2002). We additionally simulated 10^4^ and 10^5^ gene trees for the single hybridization scenario (Fig. 1a) when *τ*_1_ = 0.05, *τ*_2_ = 1.5, and *γ* ∈ {0, 0.1, 0.2, 0.3, 0.4, 0.5} to increase the size of datasets by a factor of ten and one hundred, respectively, to further evaluate the performance of *HyDe* when the level of ILS was very high. For the introgression scenario (Fig. 1b), we varied the height of node *u* to be 6^−1^, 6^−2^, 6^−3^, 6^−4^, and 6^−5^ of *τ*_1_ with *γ* ∈ {0, 0.1, 0.2, 0.3, 0.4, 0.5} to evaluate the behavior of *HyDe* when the timing of the introgression event varied. While *ms* only takes input in a tree format and cannot be used to simulate gene trees directly from a network, we note that a network can be decomposed into a forest of trees since *r* reticulations display up to 2^*r*^ trees that can be obtained by removing one of the two incoming hybrid edges for each hybrid node (denoted as *z* in Fig. 1a and 1b). We generally followed the parameter settings in Gerard et al. (2011) and assumed an island model with no migration between equally-sized, non-recombining, and panmictic subpopulations following divergence. We used the *ms* command `-ej t i j` that moves all lineages in subpopulation `i` to subpopulation `j` at time `t` that corresponds to the desired *τ* for each tree. Each parent population contained 10 individuals, five individuals were assigned to the hybrid or introgressant population(s), and one to the outgroup population. Note that the branch lengths were halved when constructing the command blocks as *ms* uses coalescent units of 4*N* generations rather than the 2*N* generations used for common models. Gene trees were simulated in proportion to the desired *γ*. For example, Figure 1a was decomposed into two trees and we simulated 10^3^*γ* gene trees from one tree obtained from the model network

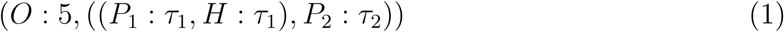

and 10^3^(1 − *γ*) from the other tree

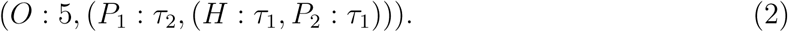

Similarly, the networks in Figure 1b–1f were broken down into 2^*r*^ trees and the number of gene trees simulated for each tree was 10^3^ *×* (2^−*r*^).

The simulated gene trees were then used as input to simulate DNA sequences using *seq-gen* (Rambaut and Grassly, 1997) with a length of 100 base pairs per gene to mimic the short-read lengths generated by next-generation sequencing methods (Kubatko and Chifman, 2019). We scaled each branch of the input gene trees by setting the parameter `-s` as 0.036, to make them equal to the expected number of substitutions per site for each branch (Rambaut and Grassly, 1997). The sequences were generated under the HKY model with *κ* = 1, and A,C,G, and T frequencies of 0.300414, 0.191363, 0.196748, and 0.311475, respectively, as described in Yu et al. (2014) and Solís-Lemus and Ané (2016). The sequences were formatted using the `concat` function in *goalign* (https://github.com/fredericlemoine/goalign) followed by single-nucleotide polymorphism (SNP) extraction and format conversion into `fasta`, `phylip`, and `vcf` for subsequent analyses using *snp-sites* (Page et al., 2016). The SNP data were converted into *structure* readable format using *PGDSpider* (Lischer and Excoffier, 2012) and ADMIXTURE readable `.bed` format using *plink2* (http://www.cog-genomics.org/plink/2.0/; Chang et al., 2015). In addition, we produced a truncated version of the simulated datasets that only contained one individual for each population as required for the computation of the *D*-statistic.

### Analysis

We used the *D*-statistic (Green et al., 2010; Durand et al., 2011; Patterson et al., 2012), *HyDe* (Blischak et al., 2018; Kubatko and Chifman, 2019), *structure* (Pritchard et al., 2000), and ADMIXTURE (Alexander et al., 2009) to detect hybridization in the simulated sequence datasets. The former two methods provide formal statistical hypothesis tests based on expectations for the site pattern frequencies in the absence of hybridization, while the latter two are commonly used population clustering methods that provide estimates of ancestry coefficients for the sampled sequences.

The *D*-statistic considers bi-allelic sites for four-taxon introgression networks (i.e., Fig. 1b). The outgroup defines the ancestral state (denoted as A) relative to the derived state (B) where A */*= B ∈ *{A, C, T, G}*, thus the model tree that can be obtained by removing the reticulation edge between nodes *u* and *z*, (((*P*_2_, *I*_1_), *P*_1_,)*O*), is supported by the site patterns BBAA or BBBA. Polyphyletic appearances of the derived state in the tree, ABBA and BABA, represent discordant site patterns, as they are not expected in the model tree mentioned above but are expected for gene trees (((*I*_1_, *P*_1_), *P*_2_), *O*) and (((*P*_2_, *P*_1_), *I*_1_), *O*), respectively. These discordant site patterns are expected to occur with equal frequency if ILS is the sole source of conflict (Goulet et al., 2017). However, introgression between *I*_1_ and either *P*_1_ or *P*_2_ will disproportionately increase the frequency of ABBA or BABA for each expected gene tree. This model assumes a single introgression event in the four-taxon network being tested and a single mutation per site with a negligible number of homoplasious substitutions. The *D*-statistic is calculated by:

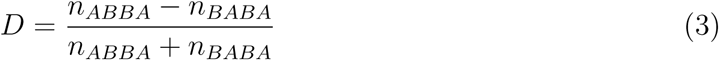

where *n*_*ABBA*_ represents the count of site pattern ABBA and *n*_*BABA*_ is the count of BABA sites.

Hybrid detection in *HyDe* is based on phylogenetic invariants, which are polynomials in the site patterns that evaluate to zero on one tree topology but are non-zero for one or more other topologies (Felsenstein, 1991; Kubatko and Chifman, 2019). Consider two tree topologies (1) and (2) obtained from Figure 1a and two linear relationships of the site pattern counts that arise on the network:

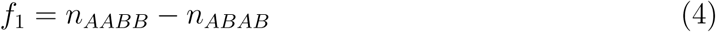

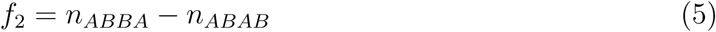

In the absence of hybridization, we expect zero when *f*_1_ is evaluated on the site-pattern counts for topology (1) and *f*_2_ for topology (2). However, neither of them is zero when the site-pattern counts are evaluated on the network with *γ* ∈ (0, 1). These functions are very attractive because the ratio of their expectations is a function of *γ*:

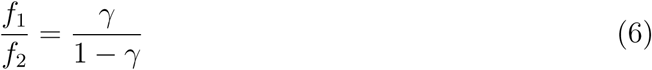

We note that this ratio is zero when *γ* = 0, which is used as the basis for the hypothesis test in *HyDe*.

Two input files are required to run *HyDe*: a multiple sequence alignment that contains at least one outgroup and three or more ingroup populations in `phylip` format, and a `map` file, a two-column text table with individual names in the first column and the name of the population to which each individual belongs in the second column. The order of the individuals in the `map` file must match the order of individuals in the `phylip` alignment. Because the order in the `phylip` file produced by our simulation pipeline changes in every replicate, the corresponding `map` file must be re-created in every run, which we automated using a custom script. The *D*-statistic was calculated on the truncated dataset using ABBA and ABAB counts reported in the *HyDe* output using a custom *R* script. We used ABAB instead of BABA because the order in which *HyDe* counts the site patterns is to list the ingroup individuals first followed by the outgroup, as opposed to the model for the *D*-statistic, which lists the outgroup first. In some cases where position of P_1_ and P_2_ were switched, particularly when *γ* = 0, we used the frequency of AABB instead of ABBA, as the tree assumed in this case would be (*O*, ((*P*_2_, *H*), *P*_1_)) instead of topology (2). While the *z*-score is often used to assess significance since *z* ⩾ 3 corresponds to *p <* 0.0013 (Zheng and Janke, 2018), an extra step to compute the *p*-value was conducted to allow objective comparisons with the *p*-value in the output of *HyDe*.

*Structure* and ADMIXTURE are model-based approaches widely used in hybridization studies to estimate ancestry coefficients for all samples in a formal statistical framework. The BI-based method *structure* infers population structure using genotype data consisting of unlinked markers by utilizing a Markov chain Monte Carlo (MCMC) algorithm to sample the relevant posterior distribution. The model assumes a user-specified number of ancestral populations in the sample (denoted by *κ*) and the individuals are probabilistically assigned to discrete populations or jointly to two or more populations if their genotypes indicate that they are admixed. The model in *structure* also assumes that every locus is in Hardy-Weinberg equilibrium and that there is linkage equilibrium within a population. The estimation procedure in ADMIXTURE is similar to *structure*, but it focuses on maximizing the likelihood rather than estimating the posterior, in an effort to enhance the speed of estimation to accommodate larger datasets.

*Structure* analyses were conducted using the admixture model. The length of the burn-in period and the number of MCMC replicates were set to 5 *×* 10^4^ and 10^5^, respectively. There is no standardized way to estimate the minimum number of iterations needed for a given dataset, and while the number of iterations may vary between studies, our choices are appropriate considering the computational intensity required for multiple replicates. Moreover, the number of iterations used for burn-in and for drawing samples from the posterior in our analysis exceeds those used in previous comparison studies, including Oliveira et al. (2015) (105 iterations for burn-in and 106 sampled iterations) and Vähä and Primmer (2005) (10^4^ iterations for burn-in and 5 *×* 10^4^ sampled iterations). We manually chose *κ* by examining the model networks (Fig. 1) as the number of distinct parental populations observed at the present time. For example, *κ* = 2 was assumed for the single hybridization scenario (Fig. 1a) as it was expected to be composed of two parental populations and one hybrid population containing about half of the genetic information from each parent when *γ* = 0.5. Likewise, we set *κ* = {2, 4, 6, 4, 2} for the scenarios in Figure 1b–1f, respectively. While there are methods to estimate the number of genetic groups that best fits the data (e.g., STRUCTURE HARVESTER; Earl and vonHoldt, 2012), we bypassed this procedure to make as ‘fair’ comparisons as possible since we provided population information for *HyDe* in the `map` file. We did not incorporate the ancestral population for the outgroup when selecting *κ* because with only one sample for the outgroup, the method may not be able to accurately estimate population allele frequencies and in the worst case, it can disrupt the final result by unnecessarily allowing more than two ancestries for the admixed populations. Nevertheless, one extra *structure* analysis for the two-hybridization scenario (Fig. 1c) with *κ* = 5 was conducted to assess the effect of allowing an extra population to accommodate the outgroup.

Lastly, ADMIXTURE analyses were conducted using the same *κ* as *structure* analyses. Because ADMIXTURE requires the input to be in binary *plink* (`.bed`) format, only bi-allelic SNPs were included in the analyses. The analyses were terminated when the log-likelihood increased by *<* 10^−4^ between iterations. All analyses were carried out using the College of Arts and Sciences’ high-performance computing cluster (Unity) at the Ohio State University. One hundred replicates were conducted for every parameter setting.

### Evaluation

The power of a statistical test refers to the probability that the test rejects the null hypothesis (*H*_0_) when the alternative hypothesis is true. Here, only the performance of *HyDe* and the *D*-statistic were compared in terms of their power to detect hybridization since *structure* and ADMIXTURE do not perform hypothesis testing. The power in *HyDe* and the *D*-statistic is defined as the proportion of the replicates for which *H*_0_: *γ* = 0 is rejected at an overall *α* = 0.05, with a Bonferroni correction applied to account for multiple testing. Specifically, *p*-values are adjusted by dividing 0.05 by the total number of triplets tested, which can be computed by 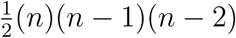 where *n* is the number of ingroup populations identified in the `map` file. The power at *γ* = 0 is equivalent to the type I error (TIE) rate. We also noted the FDR for both tests for the scenarios in Figure 1b–1f, which is the proportion of the erroneously rejected *H*_0_ in the significant observations (Benjamini and Hochberg, 1995). The FDR is calculated by:

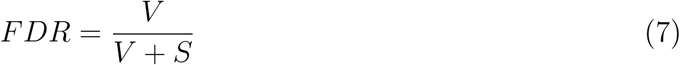

where *V* refers to the number of significant incorrect triplets and *S* is the number of significant correct triplets. The correct triplet refers to the true parent-hybrid relationship present in the model network topologies that explains a hybrid edge (Fig. 1), and the incorrect triplets refer to any other triplets. In this study, we denote a triplet by, for example, (P_1_-H-P_2_) for Figure 1a where the population H corresponds to a hybrid population and P_1_ and P_2_ represent the parents. We further relaxed the calculation of FDR for the *D*-statistic since the goal of the *D*-statistic is to detect the presence of introgression within the triplet, rather than to identify the hybrid-parent relationship. We subtracted the counts of triplets that contain at least one hybrid-parent pair, even if it only partially explains the true hybridization event, from the numerator of (7) thereby calculating a relaxed version of FDR for the *D*-statistic:

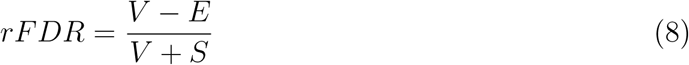

where *E* refers to the number of significant triplets that contain at least one pair of the hybrid-parent relationship. For the *structure* and ADMIXTURE analyses, we recorded the frequency with which hybrid populations were identified, noting that this measure of performance differs from the measurement of the power of a statistical test. Identifying hybrid individuals from the estimated ancestry coefficients is an inevitably subjective process; we declared an individual to be a hybrid if the estimated ancestry coefficient ∈ (0.1, 0.9).

The accuracy of the estimates of the parental contributions were evaluated by calculating the MSE of the estimated *γ* from *HyDe*, the average of the inferred ancestry of individuals (*qi*) in *structure*, and the ancestry fraction (*Q*) values in ADMIXTURE. Note that *qi* and *Q* are denoted as *γ* hereafter for convenience. The MSE is calculated by:

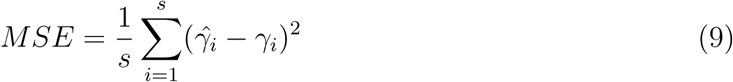

where *s* is the number of replicates (i.e., 100), 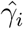 is the estimate of *γ* for dataset *i*, and *γ*_*i*_ is the true value of *γ* used to simulate the dataset *i*. The MSE can be decomposed into the sum of the variance and the square of the bias and hence low values of the MSE are indicative of estimators that are good in the sense that they minimize both variance and bias. The *D*-statistic was excluded from this comparison as it does not estimate *γ*.

Calculating the MSE was impractical for *structure* and ADMIXTURE results for the scenarios in Figure 1b–1f as more than two ancestries could be inferred for a hybrid. For these cases, we conducted Clustering Markov Packer Across K (CLUMPAK; Kopelman et al., 2015) analyses using the `LargeKGreedy` algorithm with default settings for the summation of the results. CLUMPAK was also used to eliminate the label switching phenomena in the *structure* output (Jakobsson and Rosenberg, 2007; Kopelman et al., 2015) and to reveal genuine multimodality. Multiple solutions were obtained in cases of multimodality, and we considered all distinct solutions in our analyses.

### Benchmarks

All code used to complete the simulations and analyses and all output files and R scripts to reproduce the figures and table in this manuscript are available as Jupyter Notebook and R markdown documents on Dryad (doi: ###).

## Results

### Single Hybridization Scenarios

The power to detect hybridization for *HyDe* and the *D*-statistic, as well as the frequency of hybrid identification in *structure* and ADMIXTURE, for different combinations of *τ*_1_, *τ*_2_, and *γ* in the single hybridization scenarios (Fig. 1a) are summarized in Figure 2. *HyDe* performed very well in general regardless of the number of individuals per population used, though we noted that when a single individual per population was used, both the TIE rate and the power were slightly higher than when multiple individuals were included in the sample. The *D*-statistic detected hybrids with similarly high power with negligible TIE rates for most combinations of parameters (⩽0.06, except for 0.19 when *τ*_1_ = 0.5 with *τ*_2_ = 1.5; Fig. 2a), in agreement with the results in Kubatko and Chifman (2019). *HyDe* did not perform well when ILS was prevalent when *τ*_2_ = 0.05 for all *γ* and when *τ*_2_ = 0.5 for *γ* = 0.1, where the power ranged from 0–0.1 (Fig. 2b). However, the power in *HyDe* substantially increased to ⩾0.95 when we increased the dataset size by a factor of ten, although this was accompanied by a high TIE rate (Fig. 4a). Both detection power and accuracy of the estimates of *γ* greatly improved (⩾0.97 for *γ* ⩾ 0.2) with no TIE when the size of the dataset was increased by a factor of 100 (Fig. 4b). The power for the *D*-statistic was also relatively poor (0.03–0.7) in high ILS scenarios, however, it also recovered to 1 as *τ*_2_ was increased to 0.5.

**Fig. 2.**
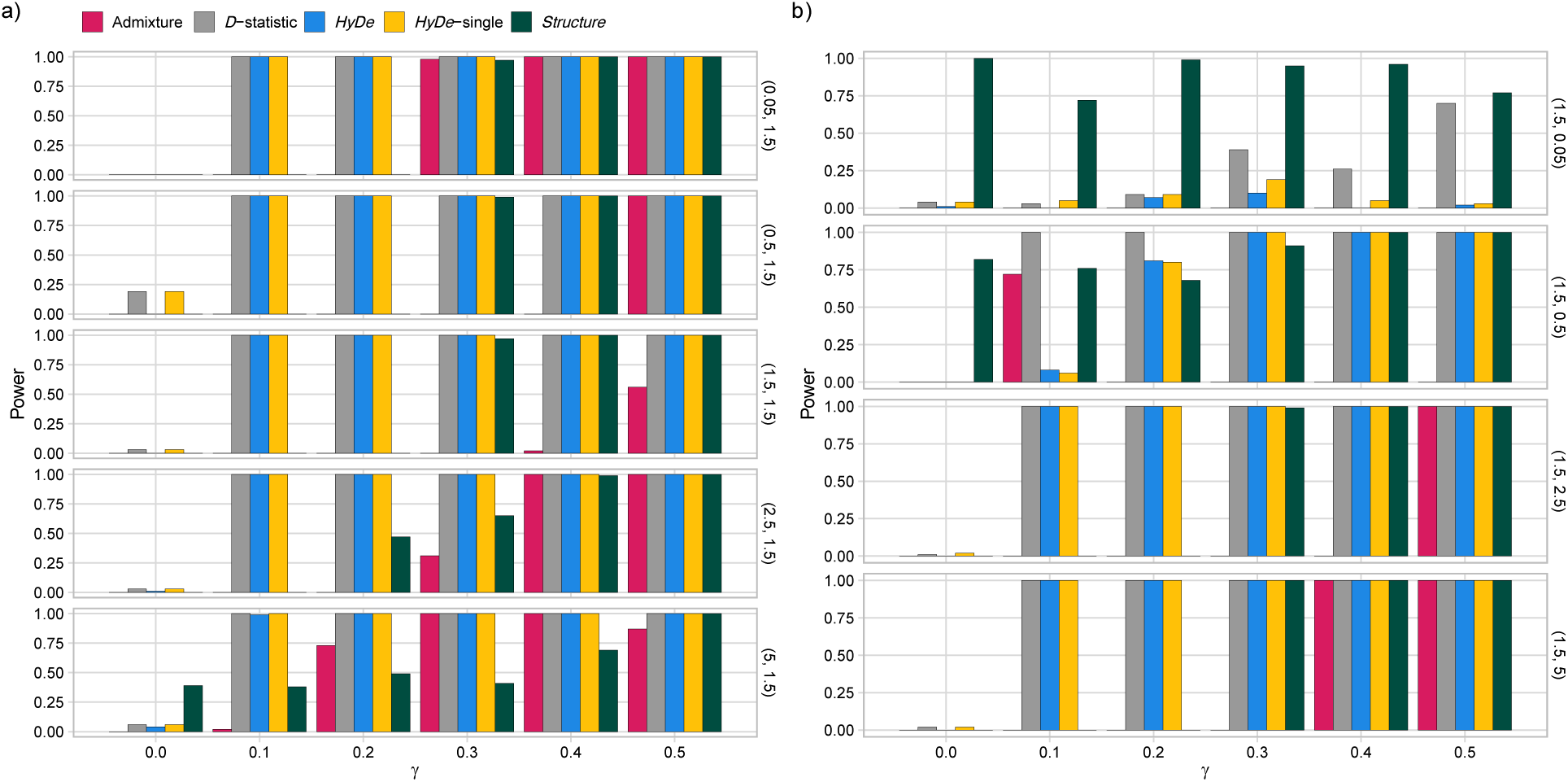
The power to detect hybridization for the *D*-statistic and *HyDe* (*HyDe*-single refers to the power when a single individual per population is used) and the frequency of hybrid identification in ADMIXTURE and *structure* for different combinations of parental contributions, *γ* ∈ {0, 0.1, 0.2, 0.3, 0.4, 0.5}, with a) *τ*_1_ ∈ {0.05, 0.5, 1.5, 2.5, 5} and *τ*_2_ fixed at 1.5; and b) *τ*_1_ fixed at 1.5 and *τ*_2_ ∈ {0.05, 0.5, 2.5, 5} in single hybridization scenarios (Fig. 1a). The two numbers in parenthesis on right side of each plot specify (*τ*_1_, *τ*_2_).

ADMIXTURE failed to identify any hybrid samples in many cases, particularly when the parental contributions were asymmetric and/or the amount of ILS was high, although one exception occurred when *τ*_2_ = 0.5 at *γ* = 0.1. Similarly, *structure* did not perform well when *γ*⩽ 0.2, with some exceptions when *τ*_1_ ⩾ 2.5 or *τ*_2_ ⩽ 0.5. *Structure* also showed high rates of false positives (0.39–1) when *τ*_1_ *<* 2.5 or *τ*_2_ ⩽ 0.5 where it detected admixed population at *γ* = 0.

Figure 3 presents estimates of *γ* and their MSE from ADMIXTURE, *HyDe*, and *structure* for different combinations of *τ*_1_, *τ*_2_, and *γ* in the single hybridization scenario (Fig. 1a). For the estimates in ADMIXTURE, the frequency with which hybrids were identified was generally the highest for the symmetric parental contributions, though the MSE was not always the smallest for *γ* = 0.5. ADMIXTURE failed to identify hybrids in most cases when *τ*_2_ was varied where the observations were not sufficient to find a pattern in the method’s performance. Similarly, even when *structure* identified hybrids when *γ* ⩽ 0.2 and *τ*_1_ ⩾ 2.5 or *τ*_2_ ⩽ 0.5, the estimates were relatively inaccurate as reflected by their high MSEs. In many cases, the *structure* estimates exhibited multimodality by clustering above and below the true *γ*, and the difference between the two clusters (i.e., modes) decreased as *τ*_1_ shortened or *τ*_2_ lengthened. While the absolute value of the *structure* estimates generally increased with *γ*, such a pattern was hard to find for high ILS scenarios. *HyDe* estimates *γ* much more accurately than both *structure* and ADMIXTURE in all cases. Even when *HyDe* performed poorly (e.g., for *τ*_2_ = 0.05), its estimates of *γ* were more accurate than the estimates from *structure*. Moreover, the *HyDe* estimates for the high ILS scenario became more accurate as the number of loci increased (Fig. 4), illustrating that the inaccurate estimates were due to inadequate signal in the data rather than methodological incompetency.

**Fig. 3.**
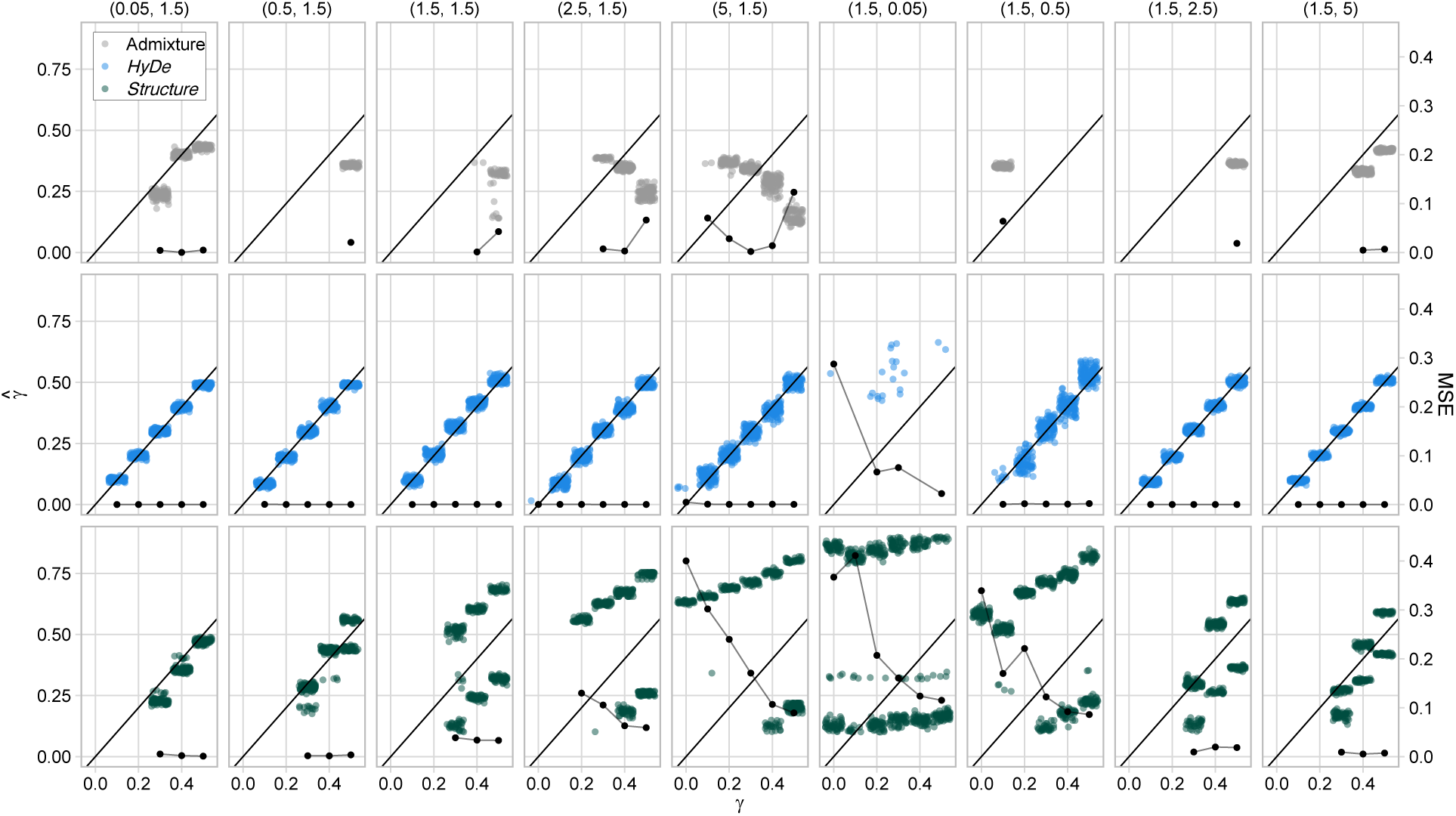
Estimates of *γ* (denoted as 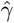 on the left y-axis) and the mean squared error (MSE; right y-axis) in ADMIXTURE, *HyDe*, and *structure* for the different combinations of parameters: *τ*_1_ ∈ {0.05, 0.5, 1.5, 2.5, 5} with *τ*_2_ fixed at 1.5 or *τ*_1_ fixed at 1.5 with *τ*_2_ ∈ {0.05, 0.5, 2.5, 5} at *γ* ∈ {0, 0.1, 0.2, 0.3, 0.4, 0.5} in single hybridization scenarios (Fig. 1a). The two numbers in parenthesis on top specify (*τ*_1_, *τ*_2_). The black solid line represents the expected *γ* when MSE = 0, the color-coded unconnected points represent the estimates of *γ* for each tool, and the connected black line with points represents the MSE.

**Fig. 4.**
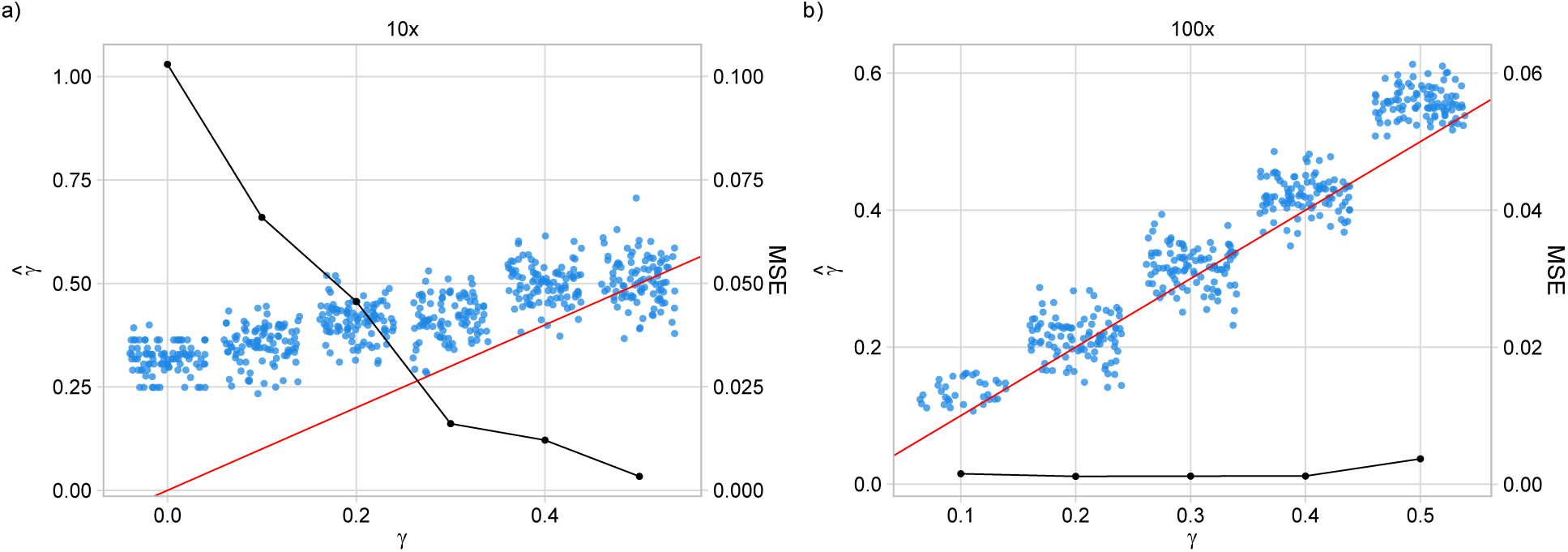
Estimates of *γ* (denoted by 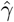 on the left y-axis) and the MSE (right y-axis) of *HyDe* for single hybridization scenarios (Fig. 1a) when *τ*_1_ = 0.05 and *τ*_2_ = 1.5 at *γ* ∈ {0, 0.1, 0.2, 0.3, 0.4, 0.5} when the dataset size was increased by a factor of a) ten (1,000 loci) and b) 100 (10,000 loci). The red solid line represents the expected *γ* when MSE = 0, the unconnected blue points represent the estimates of *γ*, and the connected black line with points represents the MSE.

### Introgression, Multi-hybridization, and Complex Hybridization Scenarios

For the introgression, multi-hybridization, and mixture of recent and ancestral hybridization scenarios (Fig. 1b–1f), both *HyDe* and the *D*-statistic exhibited a detection power of 1 for all correct triplets regardless of the number of individuals sampled from the populations. The FDR (7) for *HyDe* was generally negligible in all analyses except for the mixture of ancestral and recent hybridization events (Fig. 1f). The *D*-statistic showed very high FDR in all scenarios, ranging from 0.6 to 0.947 (Table 1), although these should be interpreted with caution. When the triplets that contain at least one pair of the hybrid and a parent were used to calculate the rFDR in (8), the rate substantially decreased, becoming as low as zero for the introgression and mixture of recent and ancestral hybridization scenarios. This is intuitive because any permutation of the populations in these scenarios will contain at least one hybrid-parent pair. In contrast, the rFDR for the multi-hybridization scenarios remained high, ranging from 0.228 to 0.316.

**Table 1.** *The false discovery rate (FDR) and the relaxed FDR (rFDR) in HyDe and the D-statistic for the introgression, multi-hybridization and complex hybridization scenarios (Fig. 1b–1f)*.

| Hybridization Scenario | FDR-*HyDe*<sup>1</sup> | FDR-*HyDe*-single<sup>2</sup> | FDR-*D* | rFDR-*D* |
| --- | --- | --- | --- | --- |
| Introgression (Fig. 1b) | 0 | 0 | 0.667 | 0 |
| Two independent events (Fig. 1c) | 0.015 | 0.043 | 0.9 | 0.3 |
| Three independent events (Fig. 1d) | 0.007 | 0.013 | 0.947 | 0.316 |
| Three hybridization events that overlap (Fig. 1e) | 0 | 0.004 | 0.69 | 0.228 |
| A mixture of ancestral and recent events (Fig. 1f) | 0.5 | 0.501 | 0.6 | 0 |

Note: <sup>1</sup>multiple individuals per population were included; <sup>2</sup>a single individual per population was included.

Figure 5a shows estimates of *γ* and their MSE in *HyDe* and *structure* in the introgression scenario (Fig. 1b). While the two methods identified introgressants in all replicates, ADMIXTURE failed to detect any. The accuracy of the estimates in *structure* was good with MSE *<* 0.015, although the estimates showed multimodality as was observed in the single hybridization scenarios. The estimates in *HyDe* were more accurate than *structure* but with slightly larger MSE (*<* 0.009) than its estimates in the single hybridization scenarios. The *HyDe* estimates and the MSE for various timings of the introgression event and for various choices of *γ* are illustrated in Figure 5b. The estimates were more accurate when introgression (node *u* in Fig. 1b) occurred later than the split event (node *v*) and when the parental contributions were more asymmetric. However, the accuracy recovered for the more symmetric parental contributions as the distance between *u* and *v* was shortened.

For the multi- and complex hybridization scenarios (Fig. 1c–1f), estimates of *γ* and the MSE in *HyDe* are shown in Figure 6. A graphical display of all distinct solutions of the ADMIXTURE and *structure* replicates identified via CLUMPAK analysis are presented in Figure 7. In the scenario with two recent hybridization events (Fig. 1c), *HyDe* unequivocally (i.e., no other triplet was found to be significant except the true ones) and accurately estimated *γ* for both hybrid edges (Fig. 6a) with MSE *<* 0.001. Assuming *κ* = 4, CLUMPAK summarized ADMIXTURE replicates into two distinct solutions (i.e., modes; Kopelman et al., 2015) where one correctly identified both hybrid populations and their parents with *γ* ≈ 0.35 (Fig. 7a; contained 61% of replicates) whereas the other solution only identified H_1_ with *γ* ≈ 0.65 (Fig. 7b; contained 39% of replicates). *Structure* replicates found three distinct solutions where two of them identified only one admixed population, either H_1_ or H_2_, with contributions from three ancestries that include both parents as well as the outgroup population in a ratio of approximately 0.38, 0.38, and 0.24, respectively (Fig. 7c and 7e; contained 68% of replicates combined); however, one solution did not find any admixed population (Fig. 7d; contained 32% of replicates). When we repeated the *structure* analysis with *κ* = 5 to prevent the outgroup contributing to the admixed population by assigning it as a distinct population, four distinct solutions were identified. Three out of the four solutions failed to identify any admixed population (i.e., ancestry coefficient *<* 0.1; Fig. 7f–7h; contained 90% of replicates combined) but one solution that was supported by 10% of replicates identified both H_1_ and H_2_, but contributions from three ancestries that included two parents and one outgroup population persisted (Fig. 7i).

**Fig. 5.**
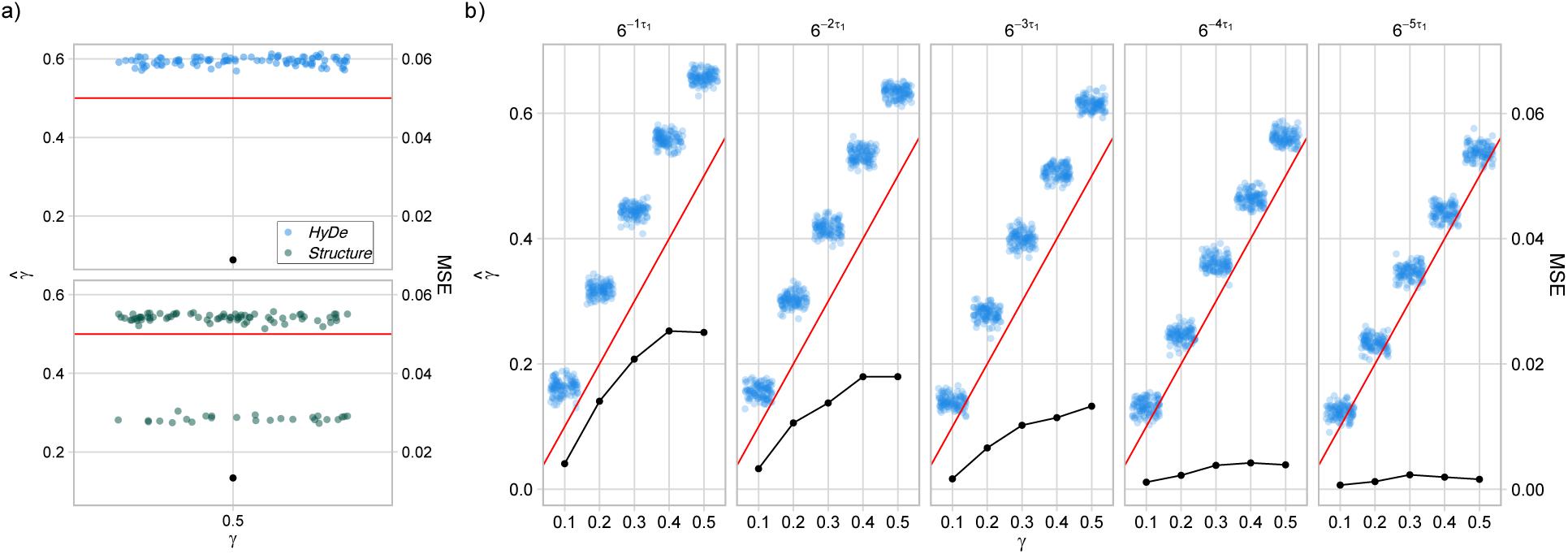
a) Estimates of *γ* and the MSE of *HyDe* (top) and *structure* (bottom) for the introgression scenario (Fig. 1b). b) Estimates of *γ* and the MSE for the introgression scenario in *HyDe* for varying timings of the introgression event. Each plot is labelled at the top by the time of introgression that represents the position of the node *u* in the model network. The time of introgression gets closer to *τ*_1_ from left to right. The connected or unconnected black points represent the MSE, and the red solid line represents the expected value of *γ* when MSE = 0.

**Fig. 6.**
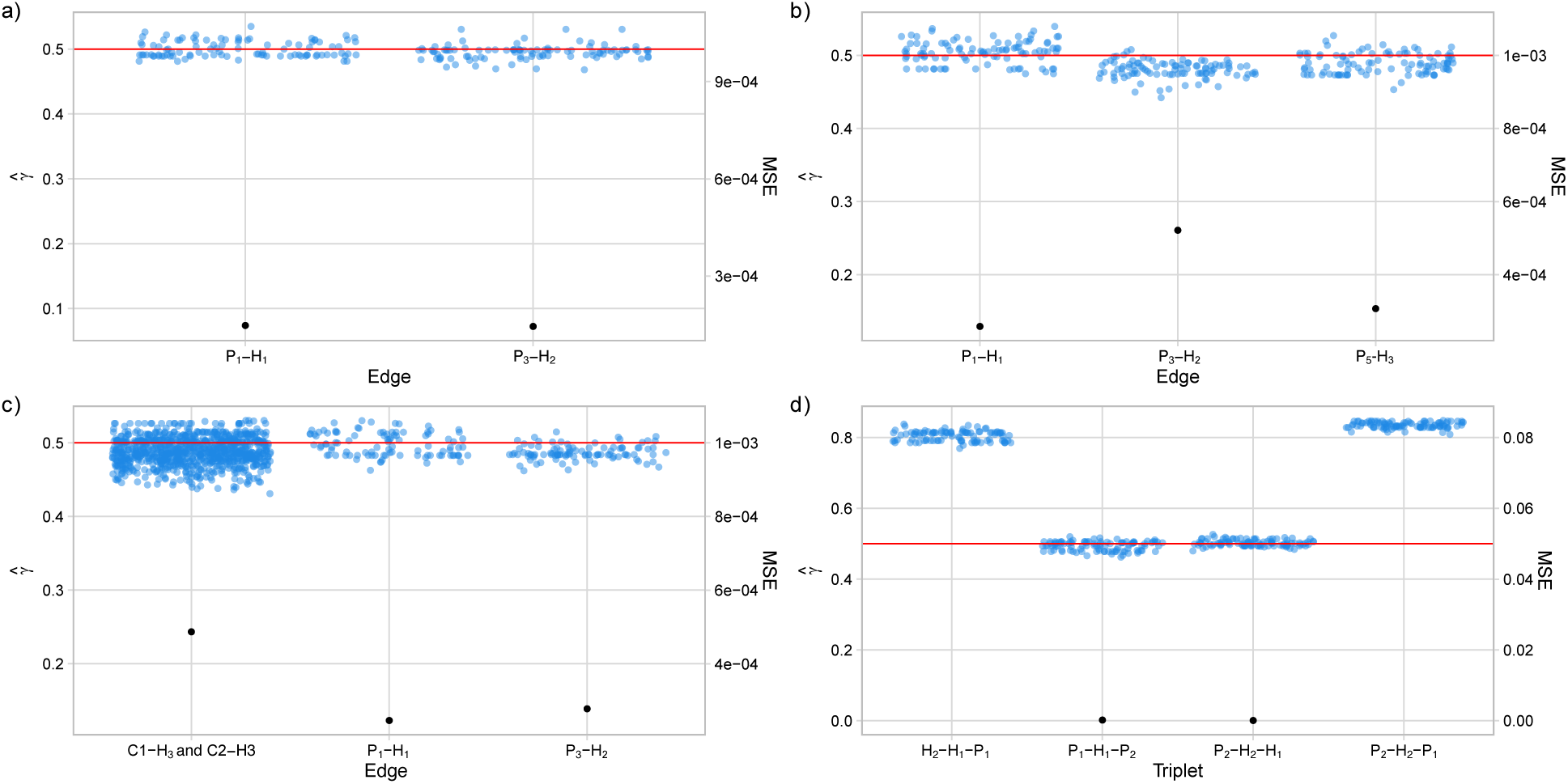
Estimates of *γ* and the MSE of *HyDe* for the a) two independent hybridization events, b) three independent hybridization events, c) three overlapping hybridization events, and d) the mixture of ancestral and recent hybridization events illustrated in Figures 1d–1f, respectively. The horizontal red line represents the true values of *γ*, and the black points represent the MSE for the corresponding hybrid edge. The blue points in a), b), and c) represent the estimates of *γ* for the corresponding edge that links the parent-hybrid pair labelled in the x-axis where C1 and C2 in c) represent any member in the clades (P_1_, H_1_, P_2_) and (P_3_, H_2_, P_4_) in Figure 1e, respectively. d) The estimates of parental contributions from the first population in the corresponding triplets labelled in the x-axis to the second population.

**Fig. 7.**
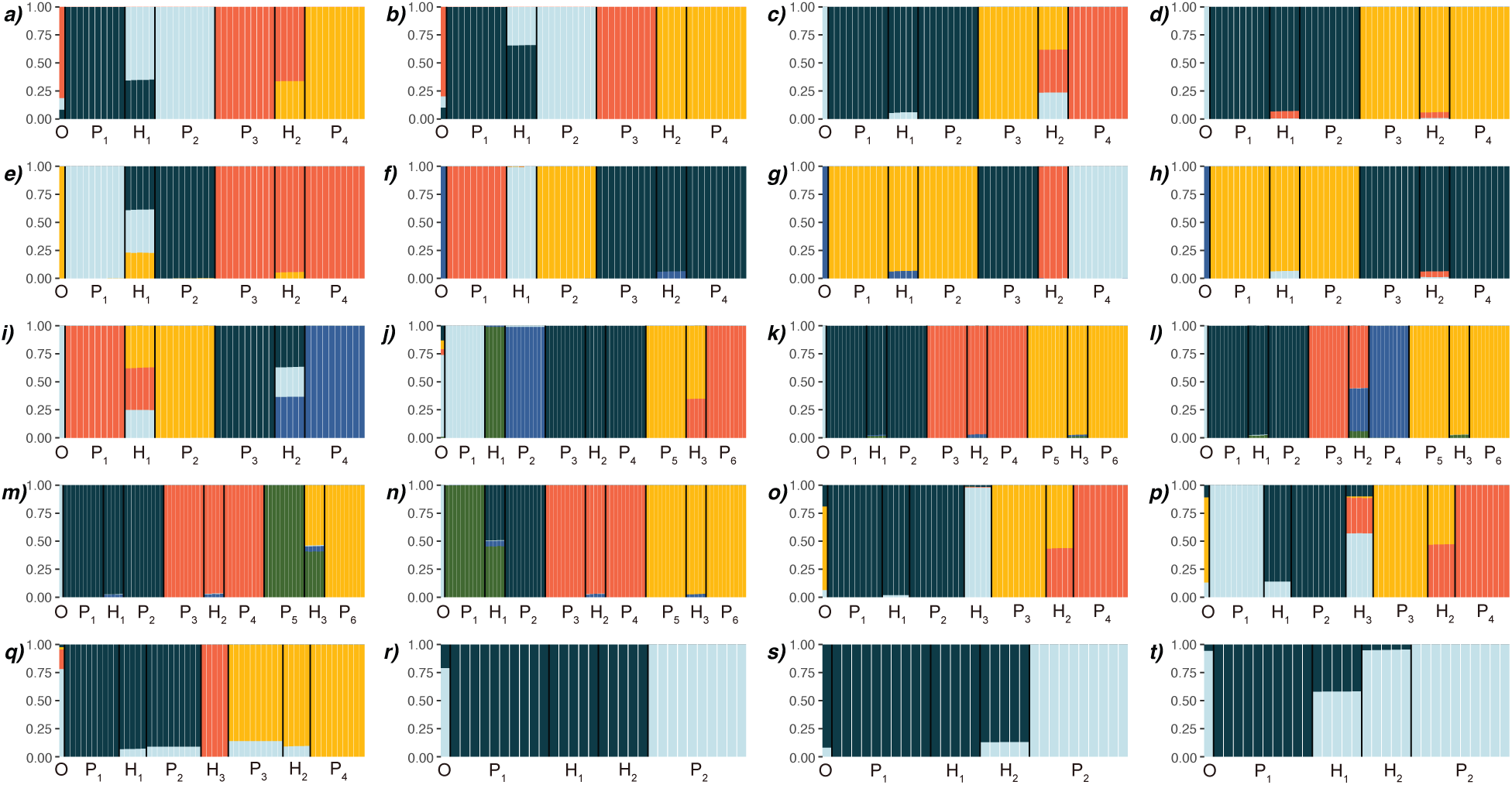
Distinct Clustering Markov Packer Across K solutions for a–b) ADMIXTURE and c–e) *structure* analyses at *κ* = 4 for the scenario with two independent hybridization events (Fig. 1c); f–i) *structure* analysis at *κ* = 5 for the scenario with two independent hybridization events (Fig. 1c); j) ADMIXTURE and k–n) *structure* analyses at *κ* = 6 for the scenario with three independent hybridization events (Fig. 1d); o–p) ADMIXTURE and q) *structure* analyses at *κ* = 4 for the scenario with three overlapping hybridization events (Fig. 1e); and r) ADMIXTURE and s–t) *structure* analyses at *κ* = 2 for the scenario with mixture of ancestral and recent hybridization (Fig. 1f). Each population is delimited by black line in the plot and labeled on x-axis; y-axis denotes ancestral coefficients.

The accuracy of *γ* estimates in *HyDe* for the scenario with three recent independent hybridization events (Fig. 1d) was outstanding, with the MSE *<* 0.001 for each of the three hybrid edges (Fig. 6b). Assuming *κ* = 6, all ADMIXTURE replicates converged into a single solution that identified only one of the three admixed populations, H_3_, with *γ* ≈ 0.35 (Fig. 7j). Four distinct solutions were identified from the *structure* replicates where one solution that contained 51% of replicates (Fig. 7k) did not identify any admixed population and each of the other three solutions identified a single admixed population (Fig. 7l–7n) that consists of four ancestries that includes both parents, the outgroup, and an ancestry that does not characterize any of the contemporary populations. The two parental populations, however, constituted the majority (*<* 0.9) for these admixed populations.

In the scenario with an ancient hybridization event followed by two recent hybridization events (Fig. 1e), *HyDe* accurately estimated *γ* with MSE *<* 0.001 for all 11 correct triplets (Fig. 6c). Note that there are nine possible permutations that explain the parent-hybrid relationships for H_3_ where one of the parents is from the clade with (P_1_, H_1_, P_2_) and the other is from the clade with (P_3_, H_2_, P_4_). The ADMIXTURE outputs yielded two equally supported solutions for which one solution only identified one admixed population with *γ* ≈ 0.44 (Fig. 7o) and the other identified all three admixed populations, H_1_ with *γ* = 0.14, H_2_ with *γ* = 0.44, and H_3_ with contributions from all four parental populations where P_1_ and P_4_ composed the majority (≈ 0.9; Fig. 7p). The *structure* replicates identified a single solution with no admixture for any of the hybrid populations but with P_3_ incorrectly identified as admixed with *γ* = 0.14 (Fig. 7q).

Finally, Figure 1f illustrates a scenario where the product of an ancestral hybridization event becomes the parent of a recent hybrid. *HyDe* accurately estimated *γ* with MSE *<* 0.001 for the two hybrid edges present in the model network. However, it also found significance for the incorrect triplets (P_1_-H_1_-H_2_) and (P_1_-H_2_-P_2_), though they are not directly observed in the network, with *γ* ≈ 0.19 and 0.16, respectively (Fig. 6d). The MSE for these unexpected triplets was not calculated as the true *γ* for these triplets was unknown. The ADMIXTURE replicates converged to a single solution that did not identify any admixed populations (Fig. 7r). The *Structure* replicates were summarized into two distinct solutions where one identified H_2_ with *γ* ≈ 0.14 but not H_1_ (Fig. 7s; contained 54% of replicates) and the other identified H_1_ with *γ* ≈ 0.42 but not H_2_ (Fig. 7t; contained 46% of replicates).

## Discussion

### On the Performance of HyDe

We have illustrated that *HyDe* is as powerful as the *D*-statistic in detecting hybridization with negligible FDR regardless of the number of individuals per population in the dataset. Also, estimates of *γ* in *HyDe* are more robust and accurate than the estimates in *structure* and ADMIXTURE in all hybridization scenarios considered in this study. *HyDe* clearly demonstrates an advantage over the *D*-statistic in that it can naturally accommodate multiple individuals per population, thereby allowing more information to be utilized in estimating site pattern probabilities leading to more accurate detection of hybridization (Blischak et al., 2018). In *HyDe*, site patterns for the population triplets (or quartet, when outgroup is included) are counted by considering all permutations of the individuals in each population. For example, the number of unique quartets for the four population (O-P_1_-H-P_2_) would be number of individuals in outgroup 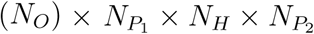 for the single hybridization scenario (Fig. 1a), and the number of site patterns for these four populations would be simply the sum of the patterns appearing in all of the permutations of individuals. In contrast, the *D*-statistic usually considers a subset of this count by including a single individual per population, for example, it only counts site patterns from a single permutation by choosing one individual from each population in the quartet (O-P_1_-H-P_2_). If the site pattern counts of multiple individuals from *HyDe* were to be directly used to calculate the *D*-statistic, then it would incorrectly find significance even in the absence of reticulation due to the amplified difference in *n*_*ABBA*_ and *n*_*BABA*_.

Nevertheless, at least three factors can impact the performance of *HyDe*. First, an excessive amount of ILS can reduce both the detection power and the accuracy of the *γ* estimates. The detection power in *HyDe* dropped from 0.99–1 to 0–0.19 (Fig. 2) when the length of *τ*_2_ was reduced to 0.05 in the single hybridization scenario (Fig. 1a). This is because the gene copy of an individual in the hybrid population, H, may fail to coalesce with the ortholog in the parent’s genome (either P_1_ or P_2_, the one that contributed the gene to H) within the very short interval between the split and hybridization event. This increases the probability that the gene copies from the two parents coalesce with each other before either coalesces with H, producing a relationship that disagrees with the model network. Consequently, the probability of each of the three possible gene tree topologies for the three ingroup populations (i.e., (*P*_2_, (*P*_1_, *H*)); (*P*_1_, (*P*_2_, *H*)); and (*H*, (*P*_1_, *P*_2_))) will be roughly one-third. In short, ILS adds to both ABBA and BABA counts, diluting information for *HyDe* and masking the footprint of hybridization within the hybrid’s genome. As expected, the power of the method increases with the sample size (Fig. 4), reconfirming that large datasets may be necessary for the optimal performance for *HyDe* when there is a high level of ILS.

Second, ancient hybridization (more precisely, an increase in the ratio of *τ*_1_ to *τ*_2_) can reduce the accuracy of the *γ* estimates. Assuming the substitution rate is homogeneous across the tree, it is intuitive that taxa with longer external branch lengths will have more time to accumulate mutations and the site patterns that would otherwise be informative for hybrid detection can be lost. Fortunately, our results indicate that the effect of increased *τ*_1_ on the estimates is not a major concern, as the largest MSE at *τ*_1_ = 5 was *<* 0.001 (Fig. 3). However, it is important to note that ancient hybridization is not uncommon in empirical systems (e.g., Genner and Turner, 2012; Escudero et al., 2014; Sun et al., 2015; Li et al., 2016; Schumer et al., 2016) and it is likely that estimating *γ* from the sampled genome would be more complex than from our simulated data. Even if the mutation rate for these populations were small, populations with longer external branches are particularly challenging due to their unique evolutionary histories during this time, including factors such as ongoing gene flow shortly after the divergence, distinct histories of introgression, maintenance of assortative mating, and rate heterogeneity (Sankararaman et al., 2014; Cahill et al., 2015).

Third, we argue that for scenarios involving introgression (Fig. 1b), as opposed to hybrid speciation (Fig. 1a), the time and direction of introgression must be taken into account for the accurate estimation of *γ*. The hybridization model assumed in *HyDe* is time-consistent (i.e., both parents of the hybrid node must coexist in time; Cardona et al., 2009), an assumption that is violated in cases of introgressive hybridization (see Fig. 1b; the vertical position of nodes *u* and *v* are different). The introgression scenario in Figure 1b does not represent direct hybridization between *u* and *v* at *z*, but illustrates our best inference of evolutionary history based on the data in present time. Based on the network, we can postulate the product of hybridization between *u* and the ancestor of I_1_ that coexisted with *u* is not extant anymore but is completely merged with I_1_ via backcrossing. In this case, because alleles in I_1_ contributed by P_1_ experienced less time to mutate than the alleles shared with P_2_, the site pattern ABBA is more likely to persist than AABB, thereby inhibiting accurate estimation of *γ* (recall (4), (5), and (6)), as evident in the larger than expected 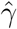 values in Figure 5b.

Moreover, a biological system can sometimes harbor both parents, a hybrid, and a backcrossing population whose evolutionary relationships can be illustrated as in Figure 1f (e.g., Wolfe et al., 1998). This scenario would also be what we might observe if the hybrid between *u* and the ancestor of I_1_ that coexisted with *u* did not go extinct and continued to backcross with I_1_. In this system, complete sampling can unintentionally complicate the interpretation of *HyDe* results because each hybrid population can have two significant parent-hybrid relationships that are not mutually exclusive. For example, in Figure 1f, emergence of H_1_ can be correctly explained by hybridization between P_1_ and P_2_ with *γ* = 0.5 or incorrectly between H_2_ and P_1_ with approximately 80% genomic contribution from H_2_. Similarly, H_2_ can have either P_2_ and H_1_ or P_2_ and P_1_ as parents also with *γ* ≈ 0.8 for the hybrid edge between H_2_ and P_2_ (Fig. 6d). It would be impossible to identify the correct triplet if the true relationship is unknown, and such significance for incorrect triplets could result in a misinterpretation of the evolutionary history of these taxa.

Our observations imply the need for a method that distinguishes hybrid speciation from introgression given the information on the site pattern frequencies. In fact, the idea of phylogenetic invariants hints at a potentially promising method. For example, note that Figure 1a can be decomposed into topologies (1) and (2) and that we will get the exact same topologies even when the positions of parents are switched. Then, the probability of the site pattern frequencies for the triplets when computed for (P_1_-H-P_2_) or (P_2_-H-P_1_) should be equal. On the other hand, if the same procedure is done for Figure 1b, the site pattern frequencies counted for the two triplets will not be expected to be equal because the two decomposed trees will have the same topology but with different branch lengths. While this idea should work well when *γ* = 0.5 and the time of introgression is sufficiently distant from the speciation event, it might not be as powerful as expected when the parental contribution is not symmetric. Progress toward a similar goal has recently been made. Hibbins and Hahn (2019) made an important contribution by proposing statistical models that can be used to test hypotheses relating to the timing and the direction of introgression that potentially allow one to distinguish hybrid speciation from introgression, although application of the method may be premature since it involves strong assumptions that may not hold for empirical datasets.

### On the Use of Population Clustering Methods for Hybrid Detection

Model-based clustering algorithms have been extremely useful in population genetic studies. Running on datasets consisting of a few loci to those with thousands of genetic markers, these methods have been shown to be powerful in classifying individual genotypes into subpopulations, a biological level at which conventional phylogenetic methods often fail to identify a resolved structure among individuals. The probabilistic quantities estimated for every individual – the ancestry coefficients – are often evolutionarily interpreted as the proportion of that individual’s alleles that are inherited from a specific ancestral population.

It is important to stress that the identification of hybrids based on the ancestry coefficient is prone to subjectivity. In practice, hybrid individuals are identified by setting parametric boundaries for the ‘pure’ populations (i.e., non-admixed populations). It is common practice to select these thresholds arbitrarily; for example, many studies define the pure population by a collection of individuals with ancestry coefficient greater than 0.8 (or 0.9) for a single ancestral population while some studies use a higher cut-off (e.g., 0.95 or even 0.99; Steeves et al., 2010). Lowering the threshold for the non-admixed populations is often justified in order to prevent overestimating the number of hybrid samples. This is a paradoxical argument because it will inevitably lead to the overestimation of the number of pure populations, particularly in case of skewed parental contributions. Sometimes the thresholds are decided *post hoc* in order to produce a desired population structure that incorporates the previous identification of individuals based on other criteria (e.g., morphology). More often, no clear justification for the choice of threshold is provided.

Even if the thresholds were appropriately justified, an individual with an estimated ancestry coefficient that is intermediate between two pure populations may not necessarily have arisen via historical hybridization. An intermediate ancestry coefficient could be observed even in the absence of hybridization when there is a large genetic divergence between populations or an excessive amount of ILS in the dataset. The former case, for instance, is observed when *τ*_1_ = 5 in Figure 1a where *structure* identified population H as admixed at *γ* = 0 (Fig. 3). In this case, even though the populations H and P_2_ share the MRCA, and are expected to have originated from the same ancestral genetic pool, the length of *τ*_1_ causes the two populations to become differentiated enough that the method cannot confidently cluster them together. However, this is less likely to be an issue in practice if *κ* can be statistically estimated as three, instead of two, as assumed in our analyses. The latter case is more problematic, because ILS and hybridization may produce similar footprint in an organism’s genome and are difficult to distinguish unless variation in the evolutionary histories of the underlying genes is taken into account, a process which is not incorporated in the models underlying the population clustering methods (Carstens et al., 2013). For example, the ancestry coefficients estimated in *structure* for the population H when *τ*_2_ = 0.05 in Figure 1a at *γ* = 0 were not distinguishable from other cases with same length of *τ*_2_ when *γ* ≠ 0. This result also illustrates that the excessive amount of ILS can override signals of hybridization in *structure* analysis. The inability to distinguish ILS from hybridization is a crucial limitation for use of population assignment methods in hybridization studies, as hybridization often occurs within closely related, rapidly evolving systems where pre- or post-zygotic barriers are not yet fully established.

Moreover, the absence of admixed populations is not evidence of the absence of hybridization for the population clustering methods. Both ADMIXTURE and *structure* often failed to identify hybrids when the parental contribution was asymmetric, while the severity varied depending on the length of *τ*_1_ or *τ*_2_. An asymmetric contribution from the parents to the hybrid is not uncommon in nature, as exemplified by *Heliconius heurippa* where the genome contains parental contributions in the ratio of 0.825 to 0.125 (Mavárez et al., 2006). Additional examples of asymmetric hybridization can be easily found in plants (Tiffin et al., 2001), invertebrates (Coyne and Orr, 1989; Presgraves, 2002), and vertebrates (Malone and Fontenot, 2008). In theses cases, hybrids are likely to be missed by the population clustering methods. Even with symmetric parental contributions, ADMIXTURE completely failed to identify introgressants and both ADMIXTURE and *structure* rarely identified all hybrid populations in the multi-hybridization scenarios. In these cases, historical hybridization in empirical systems will be underestimated despite efforts for comprehensive data collection.

Finally, it is difficult to use estimates of the ancestry coefficient to infer the hybrid status of an individual with certainty. In some studies, the numerical values of the ancestry coefficients were used to categorize admixed individuals into F_1_, F_2_, or introgressant populations. Detailed interpretation of ancestry coefficients is not recommended, not only because they may not be accurate, as evident in the high MSE in our simulations, but also because different processes can result in identical ancestry coefficients. For example, Lawson et al. (2018) explored an issue in ADMIXTURE and *structure* that produces indistinguishable plots in both the presence or absence of admixture when recent genetic drift is strong or the process deviates in other ways from the underlying inference model. Furthermore, estimates of the ancestry coefficients can result in multiple distinct solutions even when the same dataset is used, known as genuine multimodality, particularly among methods that involve stochastic simulation in the algorithm such as *structure* (Jakobsson and Rosenberg, 2007; Kopelman et al., 2015). Genuine multimodality was prevalent in our *structure* estimates, with replicates often forming two or more clusters in a single hybridization scenario and multiple sets of solutions in more complex scenarios.

Nevertheless, population clustering methods can be extremely useful in detecting hybrid populations if employed in conjunction with *HyDe. HyDe* requires information on the assignment of individuals into discrete populations, however, such information may not be always readily available for empirical datasets. In this case, population clustering analysis prior to running *HyDe* can provide important information on the number of genetically distinct populations (although accurate estimation of *κ* would still be crucial) and assignments of individuals into those populations. Moreover, results from the population clustering methods and *HyDe* may be complementary to each other in hypothesizing historical evolutionary processes. For example, a dataset may contain a high amount of ILS if an admixed population was found by the clustering methods but none of the triplets found significance for hybridization in *HyDe*, or a population may experienced multiple introgression events that result in asymmetric genomic contributions from the parents if significance was observed using *HyDe* but no admixed samples were identified in the clustering analyses.

## Conclusions

Interspecific hybridization plays an important role in nature by promoting or preventing evolutionary divergence between organisms and is being widely recognized as an essential process to understand the evolution of taxa across the Tree of Life. While accurate detection of hybrids is also a crucial step in practice (for instance, in conservation management), our results suggest that the popularly employed population clustering methods may not be reliable for such task and may potentially result in inappropriate conclusions about natural processes. Alternatively, we observe promising potential for site pattern based methods for hybrid detection that outcompete the population clustering methods in terms of both power and accuracy of their estimates of *γ*, although they may require large-scale genomic datasets to perform optimally under complex evolutionary scenarios. Finally, this study implies the need for phylogenetic network inference methods that can overcome scalability issues to simultaneously detect hybrids as well as their evolutionary history.

## Acknowledgements

We would like to thank Michael Broe and Benjamin Stone for helpful advice in developing scripts, Santiago J. Sánchez-Pacheco for useful comments that improved the manuscript, as well as Jing Peng and Kristina Wicke for discussions that refined the ideas described in this manuscript.

